# The importance of the immunodominant CD8+ T cell epitope of *Plasmodium berghei* circumsporozoite protein in parasite- and vaccine-induced protection

**DOI:** 10.1101/2020.04.03.024539

**Authors:** Matthew P. Gibbins, Katja Müller, Maya Glover, Jasmine Liu, Elyzana D. Putrianti, Karolis Bauza, Arturo Reyes-Sandoval, Kai Matuschewski, Olivier Silvie, Julius Clemence R. Hafalla

## Abstract

The circumsporozoite protein (CSP) builds up the surface coat of sporozoites and is the leading malaria pre-erythrocytic-stage vaccine candidate. CSP has been shown to induce robust CD8+ T cell responses that are capable of eliminating developing parasites in hepatocytes resulting in protective immunity. In this study, we characterised the importance of the immunodominant CSP-derived epitope, SYIPSAEKI, of *Plasmodium berghei* in both sporozoite- and vaccine-induced protection in murine infection models. In BALB/c mice, where SYIPSAEKI is efficiently presented in the context of the major histocompatibility complex class I (MHC-I) molecule H-2-K^d^, we established that epitope-specific CD8+ T cell responses contribute to parasite killing following sporozoite immunisation. Yet, sterile protection was achieved in the absence of this epitope substantiating the concept that other antigens can be sufficient for parasite-induced protective immunity. Furthermore, we demonstrated that SYIPSAEKI-specific CD8+ T cell responses elicited by viral-vectored CSP-expressing vaccines effectively targeted parasites in hepatocytes. The resulting sterile protection strictly relied on the expression of SYIPSAEKI. In C57BL/6 mice, which are unable to present the immunodominant epitope, CSP-based vaccines did not confer complete protection, despite the induction of high levels of CSP-specific antibodies. These findings underscore the significance of CSP in protection against malaria pre-erythrocytic stages and demonstrate that a significant proportion of the protection against the parasite is mediated by CD8+ T cells specific for the immunodominant CSP-derived epitope.

## INTRODUCTION

Malaria is caused by a protozoan parasite of the genus *Plasmodium* and remains a major global health challenge in tropical and subtropical countries (1). A vaccine that diminishes the burden of disease and prevents malaria transmission remains a decisive goal for malaria elimination programmes. As a gold standard in malaria vaccination, multiple immunisations of γ-radiation-attenuated *Plasmodium* sporozoites (RAS) can completely protect against wild-type (WT) sporozoite challenge (2-4). This parasite-induced protection targets the developing exo-erythrocytic forms in hepatocytes, also called liver stages, and completely abrogates blood stage infection. Antibodies and T cells have been implicated as important mechanisms of protection (5), and CD8+ T cells are the prime mediators of cell-mediated protective immunity, as exemplified in murine (6, 7) and non-human primate (8) infection models.

The circumsporozoite protein (CSP), the major surface coat protein of the malaria sporozoite, has been at the forefront of vaccination studies – being the basis of RTS,S/AS01, the most progressed malaria vaccine candidate to date (9). Immunisation of BALB/c mice with *Plasmodium berghei* (*Pb*) or *P. yoelii* (*Py*) RAS evokes immunodominant major histocompatibility complex class I (MHC-I) H-2-K^d^-restricted CD8^+^ T cell responses against distinct CSP epitopes: SYIPSAEKI for *Pb* (10) and SYVPSAEQI for *Py* (11). Indeed, the measurement of responses to these epitopes has become the standard in fundamental immunological studies in BALB/c mice (12-14). Furthermore, numerous vaccination studies involving different viral-vectored CSP- or CSP epitope-expressing vaccines – used alone or in combination as part of prime-boost regimens – have corroborated that CSP is a highly protective antigen in the BALB/c infection model (12-18). In these studies, elevated levels of either SYIPSAEKI- or SYVPSAEQI-specific CD8+ T cell responses correlated with protection.

Several studies have interrogated and contested the immunological relevance of CSP in parasite-induced protection. These studies emanated from observations that in naturally exposed humans T cell responses to CSP are scarce (19). In murine malaria models, multiple immunisations are required to elicit CD8+ T cell-dependent protective immunity in various mouse strains, particularly where no other CSP-derived CD8+ T cell epitopes have been identified (20). Furthermore, in *Py*CSP-transgenic BALB/c mice that are tolerant to *Py*CSP, complete protection can be achieved by *Py* RAS immunisation (21). In good agreement, BALB/c mice immunised with *Pb* WT parasites are completely protected when challenged with transgenic *Pb* parasites where the endogenous CSP has been swapped with the *P. falciparum* CSP (22). Taken together, these studies indicate that immune responses to CSP are dispensable for protection, and that other antigens are important to elicit protective immunity.

In this study, we have extended previous work on the entire CSP by dissecting the relevance of a single CSP-derived immunodominant epitope in parasite- and vaccine-induced protection. As the most stringent model system, we utilised transgenic *Pb* parasites lacking SYIPSAEKI for immunisation and challenge experiments in BALB/c mice. In addition, we have highlighted the level of protection achieved by CSP-based vaccines in mice expressing the relevant (BALB/c) or irrelevant (C57BL/6) MHC-I needed to present the CSP-derived immunodominant epitope.

## RESULTS

### Sporozoite-induced SYIPSAEKI-specific CD8+ T cell responses contribute to parasite killing but are dispensable for the development of sterile immunity

First, we interrogated the role that SYIPSAEKI, the H-2-K^d^-restricted immunodominant epitope of *Pb*CSP, plays in protective immunity induced after live attenuated sporozoite immunisation. For this purpose, *Pb*CSP^SIINFEKL^ radiation-attenuated sporozoites (RAS), where the SYPSAEKI sequence has been replaced with the H-2-K^b^-restricted epitope of ovalbumin, SIINFEKL ((23), Müller and Gibbins et al., unpublished) were used to immunise H-2-K^d^-expressing BALB/c mice. To date, there are no other reported H-2-K^d^-restricted *Pb*CSP epitopes identified. Removal of SYPSAEKI, by replacement with an irrelevant epitope, in the *Pb*CSP^SIINFEKL^ parasites allows unequivocal assignment of critical roles of this immunodominant *Pb*CSP-derived epitope in protection elicited by live sporozoite immunisations. Two weeks after immunisation, the frequencies of IFN-γ-producing SYIPSAEKI-specific CD8+ T cell responses in the spleen after gating for CD11a expression (an activation marker commonly used to identify antigen-experienced cells (24)) were measured by flow cytometry (Fig. 1A). As expected, *Pb*CSP^SIINFEKL^ RAS parasites elicited no SYIPSAEKI-specific CD8+ T cell responses in BALB/c mice.

**FIG 1.**
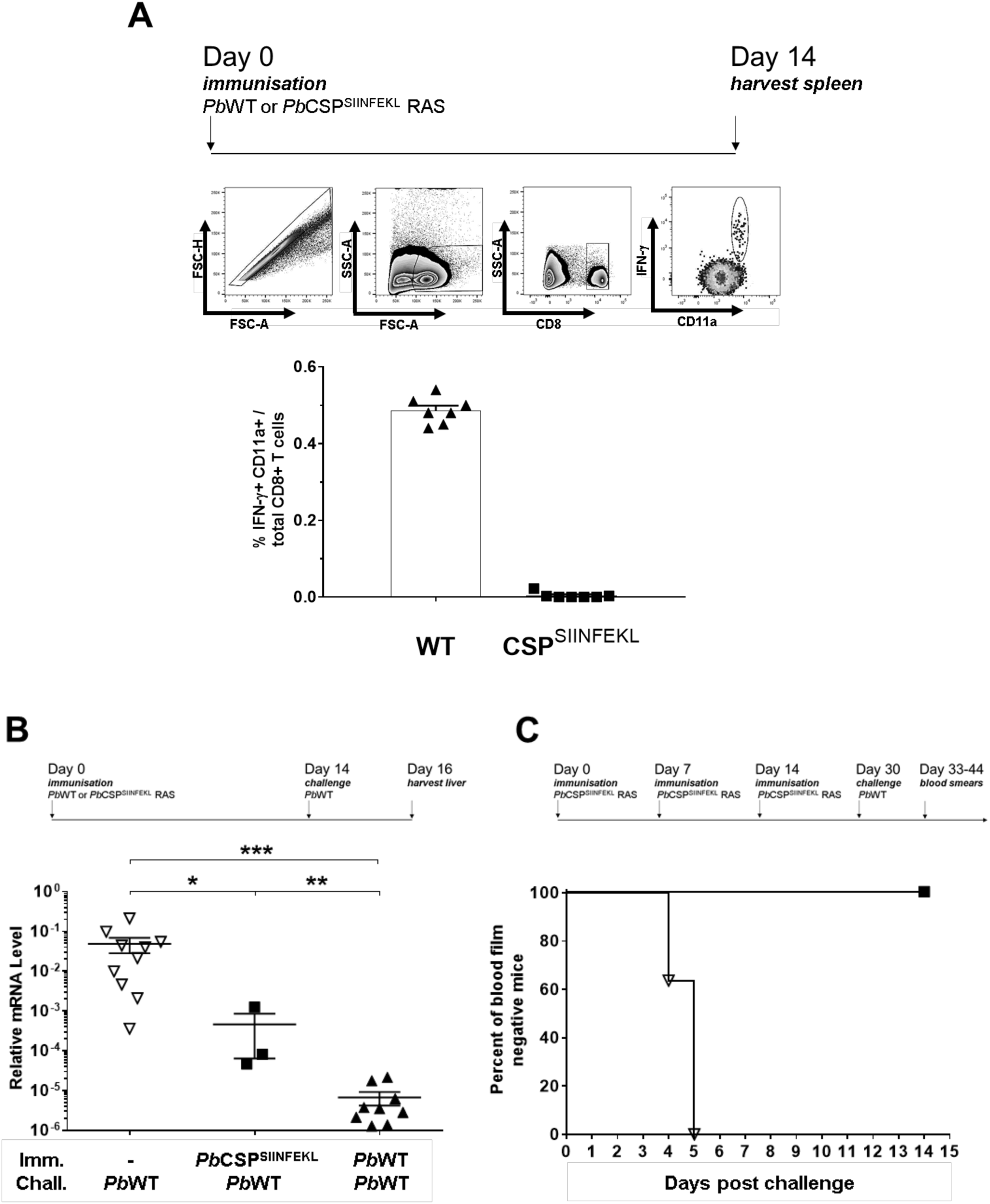
SYIPSAEKI is dispensable for RAS immunisation but predominates protection with fewer immunisations. (A) BALB/c mice were immunised once with 10,000 *Pb*WT or *Pb*CSP^SIINFEKL^ RAS. Splenocytes were taken after two weeks and restimulated with SYIPSAEKI peptide. IFN-γ-producing cells co-staining with CD11a were assessed by flow cytometry. Shown are the time course (top), the gating strategy (centre) and proportion of IFN-γ-producing CD11a of total CD8+ T cells (bottom). (B) Groups of BALB/c mice were immunised once with 15,000 *Pb*WT or *Pb*CSP^SIINFEKL^ RAS. Immunised mice and BALB/c naïve controls (n=3-10) were challenged with 10,000 *Pb*WT parasites two weeks after the last immunisation. Livers were harvested 40 hours post-challenge and the relative liver parasite loads were quantified using the ΔΔCt method comparing levels of *P. berghei* 18S rRNA and levels of mouse *GAPDH* mRNA. Mean values (±SEM) are shown and statistics were calculated using the Mann-Whitney U-test (*, p<0.05; **, p<0.01; ***, p<0.001). (C) BALB/c mice (n=12) were immunised with three doses of 10,000 *Pb*CSP^SIINFEKL^ RAS at one-week intervals. Immunised mice and naïve controls (n=11) were challenged with 5,000 *Pb*WT sporozoites 16 days after the last immunisation. Blood smears were taken daily for two weeks after challenge. Parasitaemia was assessed by microscopic examination of Giemsa-stained smears. Data shown is a combination of two independent experiments.

To ascertain whether SYIPSAEKI contributes to parasite-induced protection, BALB/c mice were immunised once with either *Pb*WT or *Pb*CSP^SIINFEKL^ RAS. Two weeks after immunisation, the mice were challenged with *Pb*WT sporozoites and protection was determined by measuring the parasite loads in the liver 40 hours later (Fig. 1B). A significant reduction in parasite load – up to four orders of magnitude difference as compared to naïve mice – was observed in mice immunised with *Pb*WT RAS and challenged with *Pb*WT parasites. In contrast, protection was reduced only by approximately two orders of magnitude in mice immunised with *Pb*CSP^SIINFEKL^ RAS (Fig. 1B). These results highlight the notion that within *Pb*CSP, the SYIPSAEKI epitope has a critical and immunodominant contribution to protecting BALB/c mice after one or two immunisations with RAS.

However, multiple immunisations with RAS are required to induce sterile protection. To establish whether the development of sterile immunity is dependent on SYIPSAEKI-specific CD8+ T cell responses, BALB/c mice were immunised thrice with *Pb*CSP^SIINFEKL^ RAS one week apart; two weeks after the last immunisation, mice were challenged with *Pb*WT sporozoites (Fig. 1C). All mice were protected from blood stage infection compared to the naïve controls, implying that SYIPSAEKI-specific CD8+ T cell responses are not necessary for the development of sterile immunity.

### Prime-boost vaccination with CSP-expressing viruses induces strong anti-CSP antibody and CD8+ T cell responses and SYIPSAEKI is the key mediator of sterile protection

Next, we probed the requirement for SYIPSAEKI presentation in protection elicited by viral-vectored CSP-expressing vaccines administered in a prime-boost regimen. Priming with adenovirus (Ad) carrying a foreign antigen and boosting with orthopoxvirus modified vaccinia Ankara (MVA) expressing the same antigen has consistently been shown to induce strong CD8+ T cell responses with high levels of protective efficacy against intracellular pathogens including malaria pre-erythrocytic stages (15, 18).

Chimpanzee adenovirus serotype 63 (AdCh63) and MVA vaccines expressing *Pb*CSP were used to vaccinate BALB/c mice with a two-week resting period between priming and boosting (Fig. 2A). Two weeks after boosting, whole blood was collected and restimulated *ex vivo* with SYIPSAEKI peptide. The frequencies of IFN-γ secreting CD8+ T cells were enumerated by flow cytometry (Fig. 2B) and Ad-MVA *Pb*CSP-vaccinated mice elicited ∼12% SYIPSAEKI-specific circulating CD8+ T cells (Fig. 2C). Serum samples were also collected from the vaccinated animals and were used in an immunofluorescence assay against air-dried *Pb* sporozoites (Fig. 2D). Ad-MVA *Pb*CSP-vaccinated BALB/c mice induced high anti-CSP antibody tires (1:10^4^). These data indicate that Ad-MVA *Pb*CSP vaccination elicit both high frequencies of SYPSAEKI-specific CD8+ T cells and high titres of CSP-specific antibodies.

**FIG. 2.**
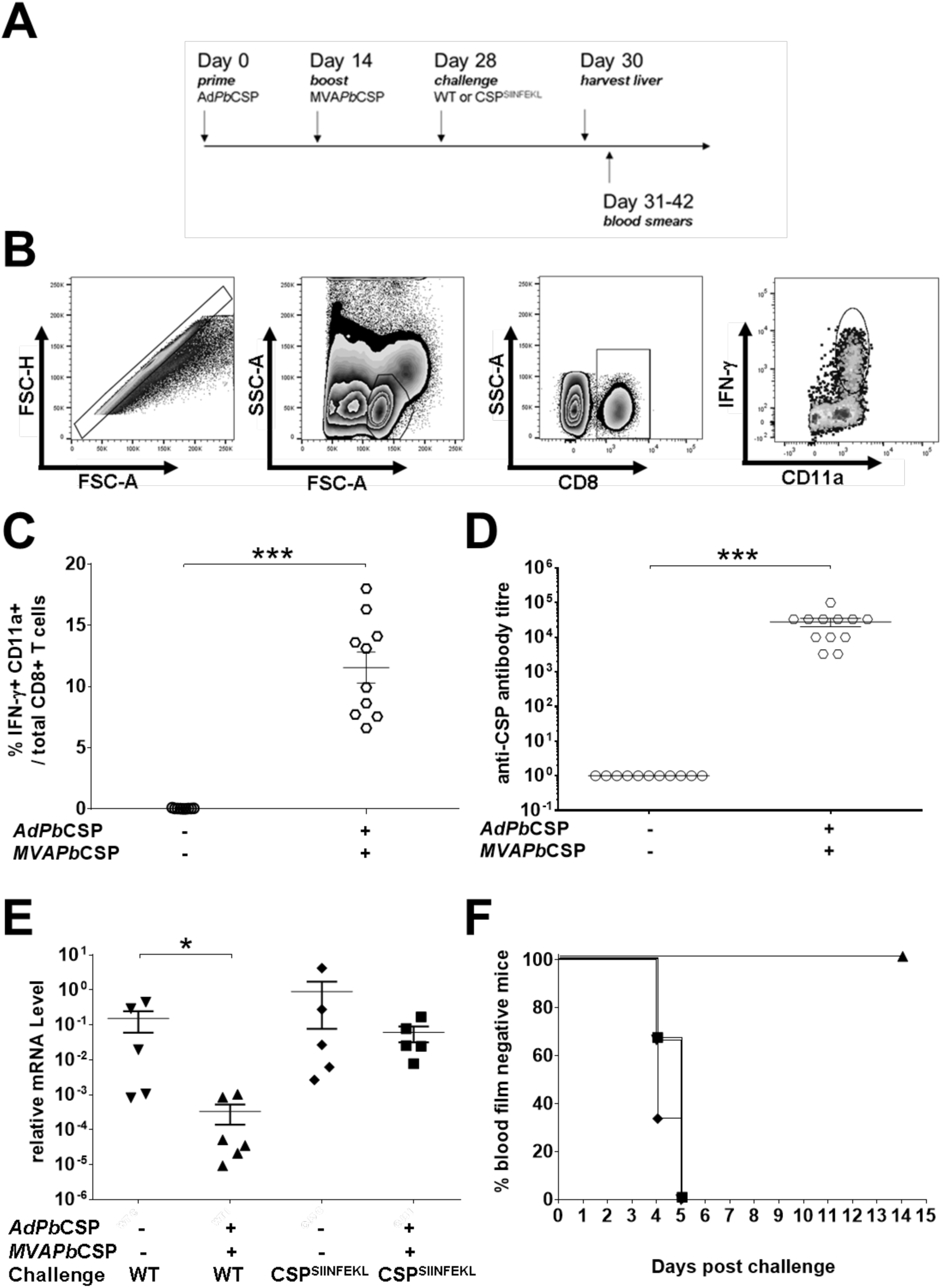
Prime-boost vaccination with viral vectored CSP-expressing vaccines induces strong anti-CSP antibody and CD8+ T cell responses, and SYIPSAEKI-specific CD8+ T cell responses are essential for protection. (A) BALB/c mice were vaccinated with AdCh63 and MVA vaccines expressing *Pb*CSP (Ad*Pb*CSP and MVA*Pb*CSP) and challenged with 10,000 *Pb*WT or *Pb*CSP^SIINFEKL^ sporozoites as shown. (B) Flow cytometry gating strategy used to determine proportions of IFN-γ+ CD11a+ CD8+ T cells. (C) Proportion of IFN-γ-producing CD11a of total CD8+ T cells. Blood was drawn from the tail from naïve (n=9) and vaccinated mice (n=10) two weeks after boost and restimulated with SYIPSAEKI and stained for CD8 and CD11a surface markers, and IFN-γ for flow cytometric analysis. (D) Reciprocal antibody titers of mouse serum reactive to whole sporozoites. Serum from naïve (n=11) and vaccinated mice (n=12) was isolated two weeks after boost and CSP specific antibody titres were measured by immunofluorescent antibody assay. (E) Livers from vaccinated mice (+) challenged with *Pb*WT (n=6) or *Pb*CSP^SIINFEKL^ sporozoites (n=5) and non-vaccinated mice (-) challenged with *Pb*WT (n=5) or *Pb*CSP^SIINFEKL^ sporozoites (n=5) were harvested 42 hours post-challenge and relative liver parasite levels were quantified using the ΔΔCt method comparing levels of *P. berghei* 18S rRNA and levels of mouse *GAPDH* mRNA. (F) Groups of vaccinated and non-vaccinated mice (n=6) were challenged with 5,000 *Pb*WT or *Pb*CSP^SIINFEKL^ sporozoites. Vaccinated mice challenged with *Pb*WT (triangles) or *Pb*CSP^SIINFEKL^ (squares) and non-vaccinated mice challenged with *Pb*WT (inverted triangles) or *Pb*CSP^SIINFEKL^ (diamonds) had daily tail smears taken from day 3-14 post challenge. Slides were stained with Giemsa and parasitaemia was assessed by microscopy. (C-E) Each data point represents one mouse with mean values (±SEM) shown and statistics were calculated using the Mann-Whiney test (*, p<0.05; ***, p<0.001).

Two weeks after boosting, Ad-MVA *Pb*CSP-vaccinated mice were challenged with *Pb*WT or *Pb*CSP^SIINFEKL^ parasites. Protection was assessed by two complementary assays; (i) determination of the reduction of parasite load in the liver (Fig. 2E) and (ii) induction of sterile protection (Fig. 2F). Strikingly, parasite load in the liver of Ad-MVA *Pb*CSP-vaccinated mice was not significantly reduced compared to non-vaccinated mice when challenged with *Pb*CSP^SIINFEKL^ sporozoites, in marked contrast to challenge with *Pb*WT sporozoites. In perfect agreement, vaccinated mice challenged with *Pb*CSP^SIINFEKL^ sporozoites were patent for parasitaemia by day 5, whereas vaccinated mice challenged with *Pb*WT sporozoites remained completely protected. These results denote that vaccine-induced effector SYIPSEAKI-specific CD8+ T responses efficiently target parasites expressing the cognate epitope. Parasites lacking the SYIPSAEKI epitope are not eliminated despite high levels of CSP-specific antibodies evoked by vaccination in this experimental system.

### CSP-based vaccines do not elicit sterile immunity in C57BL/6 mice

To further investigate the requirement of SYIPSAEKI as the indispensable protective epitope of CSP, mice unable to present this epitope were vaccinated with the *Pb*CSP prime-boost regimen with an interval of two weeks between vaccines, followed by challenge with either *Pb*WT or *Pb*CSP^SIINFEKL^ parasites (Fig 3A). C57BL/6 mice were used because SYIPSAEKI is an H-2-K^d^ restricted epitope, and this mouse strain does not express the relevant MHC-I allele. Thus, SYIPSAEKI would fail to be presented by infected hepatocytes. As before, blood and serum were derived two weeks after boost. As expected, SYIPSAEKI-specific CD8+ T cells (Fig. 3B) were not detectable in Ad-MVA *Pb*CSP-vaccinated C57BL/6 mice (Fig. 3C), but strong anti-CSP antibody titres (1:10^4^) were elicited (Fig. 3D). Ad-MVA CSP-vaccinated C57BL/6 mice challenged with either *Pb*WT or *Pb*CSP^SIINFEKL^ parasites had comparable parasite load in the liver (Fig. 3E), indicative of full liver stage development in all groups. In perfect agreement, all mice from any groups (15/15) developed blood infections.

**FIG 3.**
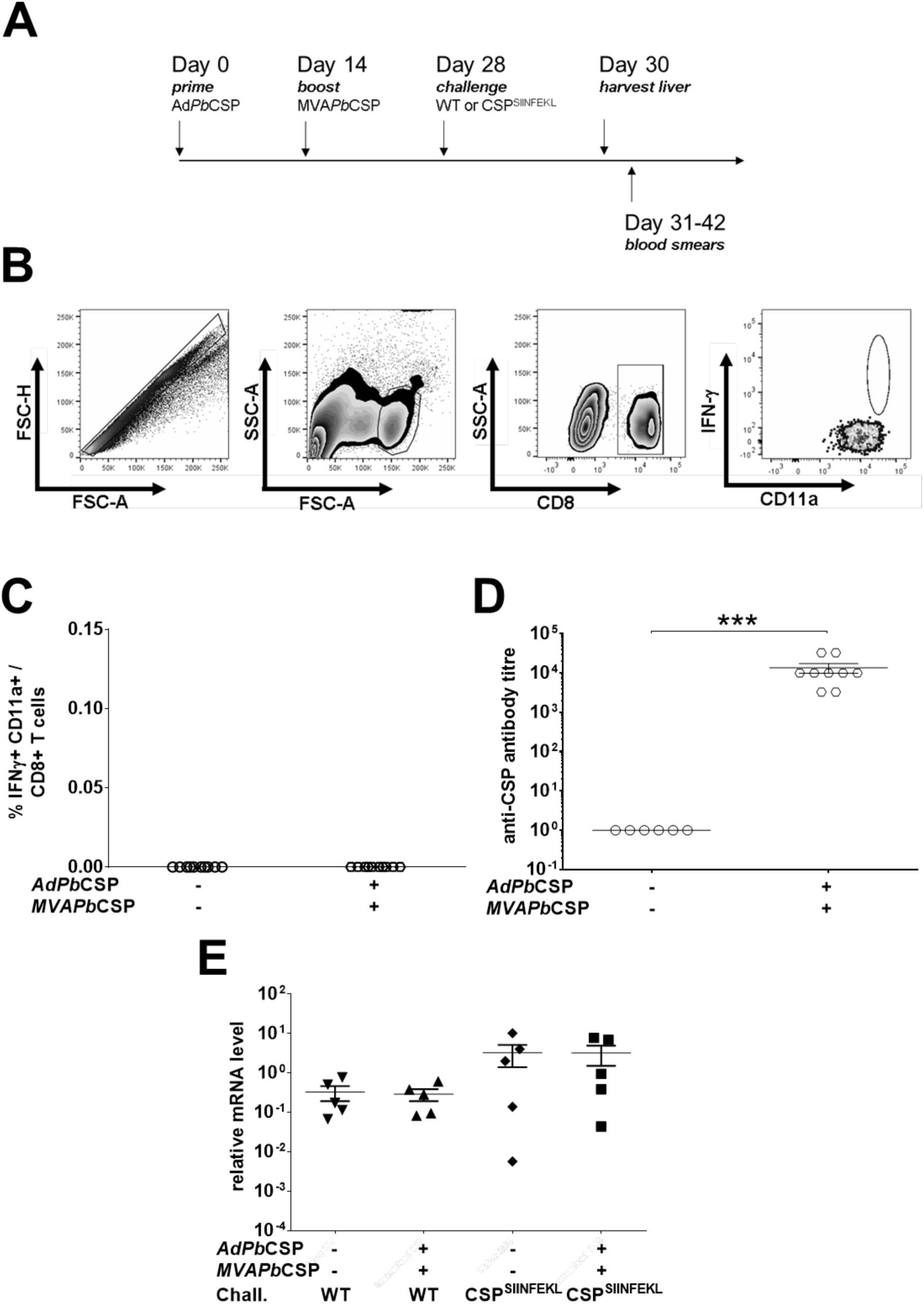
Prime-boost vaccination with CSP expressing viruses does not protect C57BL/6 mice, irrespective of induced antibody titres. (A) C57BL/6 mice were vaccinated with AdCh63 and MVA vaccines *Pb*CSP and challenged with 10,000 *Pb*WT or *Pb*CSP^SIINFEKL^ sporozoites as shown. (B) Flow cytometry gating strategy used to determine proportions of IFN-γ+ CD11a+ CD8+ T cells. (C) Proportion of IFN-γ-producing CD11a of total CD8+ T cells. Blood was drawn from the tail from naïve (n=10) and vaccinated mice (n=10) two weeks after boost was restimulated with SYIPSAEKI and stained for CD8 and CD11a surface markers, and IFN-γ for flow cytometric analysis. (D) Reciprocal antibody titres of mouse serum reactive to whole sporozoites. Serum from naïve (n=6) and vaccinated mice (n=9) was isolated two weeks after boost and CSP specific antibody titres were measured by immunofluorescent antibody assay. (E) Livers from groups of 5 mice per condition were harvested 42 hours post-challenge and relative liver parasite levels were quantified using the ΔΔCt method comparing levels of *P. berghei* 18S rRNA and levels of mouse *GAPDH* mRNA. None of the differences were significant (*p*>0.05). (C-E). Each data point represents one mouse with mean values (± SEM) shown and statistics were calculated using the Mann-Whiney test (***p<0.001).

## DISCUSSION

Our findings lend full support to the notion that CSP is an immunodominant sporozoite-derived antigen (21). A single epitope, SYIPSAEKI, is the immunodominant CD8+ T cell epitope of CSP, and we show that it is responsible for the antigen’s protective capacity against parasites in the liver in the BALB/c model. Following RAS immunisation, CD8+ T cell responses to SYIPSAEKI contribute to the reduction in parasite load in the liver following sporozoite challenge, as shown herein. When RAS-immunised mice are challenged with *Pb*CSP^SIINFEKL^, transgenic parasites lacking SYIPSAEKI, reduced anti-*Plasmodium* activity in the liver is observed. Nonetheless, complete protection is achievable in the absence of SYIPSAEKI-specific CD8+ T cell responses, demonstrating that responses to other, yet unidentified, H-2-K^d^-restricted epitopes contribute to parasite killing. It is conceivable that these epitopes are encoded by the hundreds of other *Plasmodium* genes expressed in malaria pre-erythrocytic stages, some of which might be shared with blood stage antigens (25).

Our findings also emphasise the importance of SYIPSAEKI-specific CD8+ T cell responses for promoting protective immunity when using CSP-based viral vaccines in the BALB/c model. These vaccines are aimed at generating high levels of epitope-specific memory CD8+ T cells but rely on the expression of relevant MHC-I in the vaccinated host and the presence of the cognate epitope in the parasite used for challenge (26). Notably, despite high levels of antibodies against whole sporozoites elicited following Ad-MVA *PbCSP* vaccination, sterile protection was not achieved following challenge of C57BL/6 mice. These mice cannot present SYIPSAEKI, fully supporting the notion that the protective efficacy of CSP strictly depends on the expression of the immunodominant epitope. These findings were independently corroborated by the lack of protection in mice, either BALB/c or C57BL/6, immunised with transgenic sporozoites lacking SYIPSAEKI.

Together, these results have important implications for the development of next generation malaria vaccines. We have demonstrated the significance of a single epitope of CSP in mediating protective CD8+ T cell responses while also recapitulating that protection can be achieved in the absence of responses to the entire CSP antigen (21, 22). In BALB/c mice, SYIPSAEKI-specific CD8+ T cell responses offered protection. However, to achieve complete sterile protection either multiple sporozoite immunisations or viral vaccines, which induced large populations of SYIPSAEKI-specific CD8+ T cells, were required. Multiple immunisations likely induced a broad range of immune responses and multiple high-dose immunisations with RAS in humans have been shown to induce dose-dependent anti-sporozoite CD8+ T cell responses in addition to dose dependent anti-sporozoite antibody and CD4+ T cell responses (4). In line with this, our findings lead us to suggest that future pre-erythrocytic malaria vaccine research should not only focus on inducing strong CD8+ T cell responses against one or multiple antigens, but should try to target a broad array of antigens covering diverse MHC to offer the best protection possible. The identification of novel antigens and epitopes that contribute to protection in H-2-K^d^-restricted BALB/c mice, and ultimately in human populations with broad MHC haplotypes, will aid this development. In C57BL/6 mice pre-erythrocytic immunity is mounted irrespective of CSP-specific CD8+ T cell responses, and recent genome-wide epitope profiling returned multiple sporozoite antigens and epitopes (27-29). RTS,S/AS01, the leading subunit malaria vaccine based on CSP, seems to offer some protection against *P. falciparum* re-infection (9). Partial and short-lived protection is likely primarily mediated by the action of transitory anti-sporozoite antibodies (30-32). Strikingly, peripheral blood CD8+ T cell responses were not identified to provide a role following sporozoite challenge in this candidate vaccine. Together with previous findings (7, 14, 16, 21) our data underscore efforts to improve the most advanced candidate malaria vaccine, RTS,S/AS01, by eliciting CD8+ T cells against CSP or other immunodominant antigens.

## MATERIALS AND METHODS

### Ethics and animal experimentation

Animal procedures were performed in accordance with the German ‘Tierschutzgesetz in der Fassung vom 18. Mai 2006 (BGBl. I S. 1207)’ which implements the directive 2010/6 3/EU from the European Union. The protocol was approved by the ethics committee of the Berlin state authority (‘Landesamt für Gesundheit und Soziales Berlin’, permit number G0469/09). Animal experiments at London School of Hygiene and Tropical Medicine were conducted under license from the United Kingdom Home Office under the Animals (Scientific Procedures) Act 1986. CD-1 mice were bred in-house at LSHTM, while NMRI, C57BL/6 and BALB/c laboratory mouse strains were purchased from either Charles River Laboratories (Margate, UK or Sulzfeld, Germany) or Janvier (Saint Berthevin, France). Female mice of 6-8 weeks of age were used in the experiments.

### *Plasmodium* parasites and immunization

The transgenic *P. berghei* ANKA CSP^SIINFEKL^ (*Pb*CSP^SIINFEKL^) parasite was generated with the immunodominant CSP CD8+ T cell epitope SYIPSAEKI (252-260aa) being replaced with the H-2-^b^ restricted *Gallus gallus* ovalbumin CD8+ T cell epitope SIINFEKL (258-265aa) via double homologous recombination ((23), Müller and Gibbins *et al*., unpublished). Wild-type *Plasmodium berghei* ANKA (clone c115cy1) (*Pb*WT) and *Pb*CSP^SIINFEKL^ were maintained by continuous cycling between murine hosts (NMRI or CD-1) and *Anopheles stephensi* mosquitos. Infected mosquitos were kept in incubators (Panasonic and Mytron) at 80% humidity and 20°C temperature. Sporozoites were isolated from the salivary glands and attenuated by γ-irradiation at 1.2×10^4^cGy. Mice were immunised with 10,000 sporozoites administered intravenously with multiple doses given one week apart unless otherwise stated. For challenge infections, 5,000 or 10,000 sporozoites were administered intravenously to assess sterile protection and parasite load in the liver, respectively.

### Viral-vectored CSP-expressing vaccines

AdCh63 and MVA vaccines expressing the mammalian codon-optimised fragment of *Pb*CSP were constructed and propagated based on previously published viral vectors (33, 34). The viral vectors were administered intramuscularly in endotoxin-free PBS at a concentration of 10^5^ viral particles for Ad*Pb*CSP for the prime immunisation and 10^6^ viral particles MVA*Pb*CSP for the boost immunisation.

### Immunofluorescent antibody assay

10,000 sporozoites were spotted onto epoxy coated glass slides with marked rings (Medco), dried at room temperature and stored at −20°C. Thawed slides were fixed in acetone, dried and rehydrated with PBS before incubation in 10% FCS supplemented DMEM (Gibco) for 1 hour at 37°C in a humid chamber. Serum at concentrations 1:10^3^, 1:3.3×10^3^, 1:10^4^, 1:3.3×10^4^, 1:10^5^ (and, additionally, 1:3.3×10^5^ and 1:10^6^ for C57BL/6 serum) were added to the ring wells and incubated for 1 hour at 37°C in a humid chamber. Slides were washed and stained with a mouse anti-CSP (35) primary antibody. Hoechst33342 was added as the nuclear stain together with a respective fluorescently labelled anti-mouse secondary antibody for a further one-hour incubation. Slides were washed and mounted with ‘Fluoromount-G’ (Southern Biotech) and analysed by fluorescent microscopy (Zeiss Axio Observer).

### Quantification of SYIPSAEKI-specific CD8+ T cell responses

Spleens were harvested and lymphocytes were derived by passing spleens through 40μm cell strainers (Corning). Peripheral blood was drawn from the tail vein and collected in Na^+^ heparin capillary tubes (Brand) and assayed in 96-well flat bottom plates (Corning). Red blood cells were lysed using PharmLyse (BD) and lymphocytes were resuspended in 10% FCS, 2% Penicillin-Streptomycin and 1% L-glutamine supplemented RPMI 1640 (Gibco). Splenocytes were counted using a 40x dilution with Trypan Blue (ThermoFisher Scientific) and a Neubauer ‘Improved’ haemocytometer (Biochrom). 2×10^6^ splenocytes and the lysed blood samples were prepared in 96 well plates and incubated with a final concentration of 10μg/ml of SYIPSAEKI peptide in in the presence of Brefeldin A (eBioScience) for 5-6 hours at 37°C and 5% CO_2_. For staining of cell surface markers and intracellular cytokines, cells were incubated for 1 hour at 4°C for each staining. Cells were stained for CD8 (53-6.7) and CD11a (M17/4) (eBiosience). Splenic cells were fixed with 4% paraformaldehyde and peripheral blood cells were fixed with 1% paraformaldehyde before staining for IFN-γ (XMG1.2) (eBioscience) in the presence of Perm/Wash buffer (BD) for intracellular staining. Data was acquired by flow cytometry using an LSRFortessa or LSRII (BD) and analysed using Flowjo9.5.2 (Tree Star, Inc.).

### Quantification of parasite load in the liver

Livers were harvested 40-42 hours after sporozoite challenge and total RNA was extracted following homogenisation using TRIzol (ThermoFisher Scientific). cDNA was generated using the RETROScript Kit (Ambion). Quantitative real-time PCR was performed using the StepOnePlus Real-Time PCR System and Power SYBR Green PCR Master Mix (Applied Biosystems). Relative liver parasite levels were quantified using the ΔΔCt method comparing levels of *P. berghei 18S* rRNA using specific primers and normalised to levels of mouse *GAPDH* mRNA (36).

### Assessment of parasitaemia

Sterile protection was assessed by daily blood smears, taken from mice 3-14 days after sporozoite challenge, stained with Giemsa (improved solution; VWR) to microscopically determine the presence of blood stage parasites.

### Statistical analysis

Statistical analysis was performed using GraphPad Prism v7 (GraphPad Software, Inc.). Statistics were calculated using the Mann-Whitney U test.

## AUTHOR CONTRIBUTIONS

O.S. and J.C.R.H. designed the experiments in the laboratory of K.Matuschewski; O.S. generated the transgenic parasites CSP^SIINFEKL^; M.P.G., K.Müller., M.G., J.L. and E.D.P. performed experiments and analysed data; K.B. and A.R.-S. generated the CSP-expressing viruses Ad*Pb*CSP and MVA*Pb*CSP; M.P.G. and J.C.R.H. wrote the paper. All authors commented on and approved the paper.

## ACKNOWLEDGEMENTS

J.C.R.H. was funded by grants from The Royal Society (University Research Fellowship UF0762736/UF120026 and Project Grant RG130034) and the National Centre for the Replacement, Refinement & Reduction of Animals in Research (Project Grant NC/L000601/1). O.S. was funded in part by the Laboratoire d’Excellence ParaFrap (ANR-11-LABX-0024). K.Matuschewski was supported by the Max Planck Society and grants from the European Commission (EviMalaR Network of Excellence #34), the Chica and Heinz Schaller Foundation, and the Alliance Berlin Canberra “Crossing Boundaries: Molecular Interactions in Malaria”, which is co-funded by a grant from the Deutsche Forschungsgemeinschaft (DFG) for the International Research Training Group (IRTG) 2290 and the Australian National University. A.R-S., a Jenner Investigator and an Oxford Martin Fellow, was funded by a Wellcome Trust Career Development Fellowship (Grant 097395/Z/11/Z). We would like to thank the Jenner Institute’s Viral Vector Core Facility for the viral vectored vaccines. The funders had no role in study design, data collection and analysis, decision to publish, or preparation of the manuscript.

